# Requirement of FLCN tumor suppressor gene for mTORC1-mediated inhibition of TFE3 transcriptional activity

**DOI:** 10.1101/2020.07.08.193169

**Authors:** Hua Li, HyoungBin Oh, Benoit Viollet, Laura S. Schmidt, W. Marston Linehan, Vera P. Krymskaya, Seung-Beom Hong

## Abstract

TFE3 is an oncogenic transcription factor whose subcellular localization and activity are regulated by post-translational modifications. TFE3 is hyper-phosphorylated in the cytoplasm of mammalian cells under normal growth conditions, but becomes hypo-phosphorylated and translocates into the nucleus of cells with genetic inactivation of the *FLCN* tumor suppressor gene, which is responsible for the development of renal cancer, lung cysts and cutaneous hamartomas in patients with Birt-Hogg-Dubé syndrome (BHD). Since FLCN is suggested to play a role in metabolic signaling through its interaction with energy and nutrient sensing kinases AMPK and mTOR, we investigated whether metabolic signaling regulates TFE3 activity through post-translational modifications and subcellular localization. We found that TFE3 was activated by metabolic stresses, such as glucose or amino acid deprivation, and induced gene expression of lysosomal genes, *cathepsins, V-ATPase* and *mucolipin-1*. AMPK activation and/or mTORC1 inhibition by nutrient deprivation, AMPK activator, mTOR or lysosomal inhibitors induced hypo-phosphorylation and subsequent nuclear localization of TFE3. mTORC1 directly phosphorylated TFE3 *in vitro* and the mutation of a putative mTORC1-dependent TFE3 phosphorylation site, Ser 321, was critical for its interaction with 14-3-3 protein, and retention in the cytoplasm. Direct phosphorylation of TFE3 by AMPK was demonstrated by an *in vitro* kinase assay and the AMPK-mediated phosphorylation sites were determined by mass spectrometry; however, the physiological significance of AMPK-mediated TFE3 phosphorylation requires further investigation. mTORC1-mediated TFE3 phosphorylation under nutrient and growth factor replete conditions was dependent on FLCN expression. In summary, we have shown that TFE3 induces lysosomal gene expression upon activation of AMPK and inhibition_of mTORC1 activity by metabolic stresses or loss of FLCN, which would play important roles not only in catabolic processes but also in the pathogenicity of human disease including cancer.

## Introduction

AMPK and mTORC1, both of which sense energy and nutrient levels, are master regulators of metabolism, energy homeostasis and cell growth^1,2^. Glucose deprivation reduces cellular energy levels, which in turn induces AMPK activity that suppresses anabolic processes but facilitates catabolic processes to accommodate cellular energy needs for survival. mTORC1, which senses the cellular level of amino acids, induces protein synthesis and cell growth by phosphorylating its substrates 4EBP1 and S6K1. mTORC1 activation by amino acids requires its localization to the lysosomal membrane and lysosomal V-ATPase activity^3,4^. AMPK inhibits mTORC1 by phosphorylating TSC2, a negative regulator of mTORC1, and Raptor, a major component of mTORC1^5,6^. Inhibition of mTORC1 in response to nutrient deprivation initiates autophagy, a process that requires autophagosome formation and subsequent fusion between the autophagosome and the lysosome to form the autolysosome ^4,7–10^. Cellular organelles and macromolecules are catabolized by lysosomal hydrolases in the autolysosomes to produce nutrients and energy. Inhibition of lysosomal proteinases or lysosomal V-ATPase, which maintains low pH in the lysosome, prevents degradation of autophagic substrates. Lysosomes and lysosomal hydrolases are also involved in the pathogenic processes of human disease including cancer^11^. Perturbation of autophagy contributes to tumorigenesis paradoxically by promoting both cell survival and cell death^12^.

Members of the MiTF/TFE transcription factor family (MiTF, TFE3, TFEB and TFEC) regulate the expression of lysosomal and autophagic genes^13–21^. Interestingly, gain of function of these genes is associated with the development of human cancers including malignant melanomas, clear cell sarcoma and renal cell carcinomas^22^. Accordingly, it is likely that the expression of lysosomal genes driven by MiTF/TFE transcription factors contributes to tumorigenesis and tumor progression. Gene amplification of MiTF and TFE3, and chromosomal translocations involving either TFE3 or TFEB are found in cancers and are regarded as the major mechanisms of their activation ^23–27^ However, post-translational modifications such as phosphorylation and sumoylation of MiTF, TFE3 and TFEB also play important roles in regulating their activities^14,16,18,28–30^. Several kinases including, Erk-MAPK, mTOR, C-Tak1 and GSK3 have been shown to phosphorylate MiTF/TFE transcription factors^31–34^.

Birt-Hogg-Dubé (BHD) syndrome is characterized by the development of hair follicle hamartomas, lung cysts, recurrent spontaneous pneumothorax, and renal cancer. The responsible gene *FLCN* was cloned in 2002 and was shown to have tumor suppressor function ^35–36^. FLCN forms a complex with FLCN-interacting proteins 1 and 2 (FNIP1 and FNIP2), and 5’-AMP-activated protein kinase (AMPK) and exhibits GAP activity for RagC/D that induces mammalian target of rapamycin complex 1 (mTORC1), activation^37–41^. Genetic inactivation of the *FLCN* tumor suppressor gene induced nuclear localization of TFE3, which is hyper-phosphorylated and preferentially localized to the cytoplasm in cells under normal growth conditions ^14^. Loss of FLCN induces anti-microbial genes and inhibits WNT signaling through TFEB and/or TFE3 activation^42–43^. Not only FLCN but also the functional role of AMPK in the regulation of TFEB/TFE3 axis has been demonstrated ^42,44^. Here, we sought to explore the physiological stimuli and specific kinases regulating the nucleocytoplasmic shuttling and transcriptional activity of TFE3. The current study demonstrated that metabolic stresses, such as glucose or amino acid deprivation, induced lysosomal gene expression through activation of TFE3. Involvement of FLCN, AMPK and mTORC1 in the regulation of TFE3 was investigated.

## Materials and methods

### Cells, culture conditions and reagents

Wildtype (WT) and AMPK-null (AMPK-KO) MEFs were kindly provided by Dr. Benoit Viollet (INSERM, Paris, France). Mammalian cells were maintained in DMEM supplemented with 10% FBS (HyClone) and penicillin/streptomycin (Invitrogen). For glucose deprivation, DMEM without sodium pyruvate and glucose (Invitrogen), was used with or without 10% dialyzed serum (Invitrogen) depending on each experiment. For amino acid deprivation, HBSS (Invitrogen) or RPMI lacking amino acids (US Biological) was used with or without 10% dialyzed serum depending on each experiment. Antibodies against TFE3 (HPA023881), α-tubulin, mucolipin 1 were purchased from Sigma. Antibodies against P-Raptor, Raptor, mTOR, P-AMPKalpha, AMPKalpha, P-ACC and ACC were purchased from Cell Signaling. Antibodies against Topoisomerase 1 alpha and ASAH1 were purchased from Santa Cruz. 14-3-3 antibody and active recombinant AMPK (α2β1γ1) were from Millipore. Precast Tris-Glycine polyacrylamide gels and 10X tris-glycine gel electrophoresis buffer were from BioRad. Trizol reagent, SeeBlue Plus 2 protein size marker, Superscript II reverse transcriptase, 10 mM dNTP, DNase I, and random hexamers were from Invitrogen. Power SYBR Green PCR mix was from Applied Biosystems. Cells were transfected with siRNAs or plasmid DNAs using Dharmafect 4 (Dharmacon) or Fugene 6 (Roche Applied Sciences) according to the manufacturer’s protocol. 2-deoxy-D-glucose (2-DG), Bafilomycin A1 and chloroquine diphosphate salt were from Sigma. Complete proteinase and PhosphoSTOP phosphatase inhibitor cocktails were from Roche Applied Sciences. Adenoviruses expressing wildtype FLCN and TFE3 have been described^36,45^. HA-Raptor (Plasmid 1859), myc-mTOR (Plasmid 1861), GST-14-3-3 (Plasmid 1942) expression vectors were obtained from Addgene and described ^45–46^.

### Total RNA isolation and RT-PCR

Total RNAs were isolated from cultured cells using Trizol reagent and further purified by RNeasy purification kit (Qiagen) following the manufacturer’s instructions. cDNAs were generated from the RNAs by using Superscript III reverse transcriptase and random hexamers. Quantitative RT-PCR was performed with Power SYBR-Green Master Mix using a Eppendorf Mastercycler^®^ ep realplex real-time PCR system following the manufacturer’s protocols. All reactions were run in triplicate using *β-actin*, *GAPDH* or *cyclophilin A* genes as internal controls. The gene-specific primer pairs for the PCR reactions are as follows: human actin forward 5’-ATCAAGATCATTGCTCCTCCTGAG-3’ and reverse 5’-AGCGAGGCCAGGATGGA-3’; *GAPDH* forward 5’-GGTGAAGGTCGGAGTCAACG-3’ and reverse 5’-CACCAGGGCTGCTTTTAACTCT-3’; *GPNMB* forward 5’-TGCGAGATCACCCAGAACACA-3’ and reverse 5’-CGTCCCAGACCCATTGAAGGT-3’; *Mcoln1* forward 5’-ACCCTGACATCCCCAG-3’ and reverse 5’-TTAAGGAAGCCACGGAGCAGT-3’; *Atp6v0b* forward 5’-GTCTGCGTGGGCATCGTG-3’ and reverse 5’-CCAGCAATAAATAACAGTAGGG-3’; *Gpnmb* forward 5’-CCAGCCACTTCCTCAACGA-3’ and reverse 5’-CCAGTGTTGTCCCCAAAGTTC-3’; *Actin* forward 5’-GACAGGATGCAGAAGGAGATTACTG-3’ and reverse 5’-GCTGATCCACATCTGCTGGAA-3’; *NEU1* forward 5’-CAGCACATCCAGAGTTCCGAGT-3’ and reverse 5’-TGTCTCTTTCCGCCATGAGGT-3’; *ATP6V0E1* forward 5’-CATTGTGATGAGCGTGTTCTGG-3’ and reverse 5’-AACTCCCCGGTTAGGACCCTTA-3’; *CTSA* forward 5’-CAGGCTTTGGTCTTCTCTCCA-3’ and reverse 5’-TCACGCATTCCAGGTCTTTG-3’; *CTSB* forward 5’-AGTGGAGAATGGCACACCCTA-3’ and reverse 5’-AAGAAGCCATTGTCACCCCA-3’; *CTSD* forward 5’-AACTGCTGGACATCGCTTGCT-3’ and reverse 5’-CATTCTTCACGTAGGTGCTGGA-3’; *CTSK* forward 5’-GCCAGTTTTCTTCTTGAGTTGG-3’ and reverse 5’-GGATATGTTACTCCTGTCAAAAATCA-3’; *MCOLN1* forward 5’-TTGCTCTCTGCCAGCGGTACTA-3’ and reverse 5’-GCAGTCAGTAACCACCATCGGA-3’; *MCOLN2* forward 5’-GCCCTTGTGAAAAATACCGA-3’ and reverse 5’-ATGCATCAAACACCAGGACA-3’; *MCOLN3* forward 5’-TCTGGGCTCGAGGTAGAAAA-3’ and reverse 5’-ACGGAGACATTGTATAGCTGC-3’.

### Immunoblotting

Cells were washed with cold PBS and lysed in RIPA (50 mM Tris-HCl, pH 8.0, 150 mM sodium chloride, 1.0% NP-40, 0.5% sodium deoxycholate, and 0.1% SDS) or Laemmli Sample buffer supplemented with β-mercaptoethanol, benzonase and MgCl_2_ (2 mM). Cell lysates were resolved by 4-20% or 7.5% SDS. Immunoblots were processed by the SuperSignal West Pico Chemiluminescent Detection System (Thermo Scientific) according to the manufacturer’s protocols.

### Preparation of cytoplasmic and nuclear extracts

Nuclear and cytoplasmic fractionation was performed basically following the protocol described by Andrews and Faller with minor modifications^47^. Cells were washed and resuspended in buffer A (10 mM HEPES-KOH pH7.9, 1.5 mM MgCl_2_, 10 mM KCl, 0.5 mM DTT and 0.1% NP-40) containing proteinase and phosphatase inhibitors. After 10 min of swelling, cells were vortexed for 10 sec and centrifuged for 10 sec. The supernatant was used as the cytoplasmic fraction. The pellets were resuspended in buffer C (20 mM HEPES-KOH pH 7.9, 25% glycerol, 420 mM NaCl, 1.5 mM MgCl_2_, 0.2 mM EDTA and 0.5 mM DTT, 0.1% NP-40) containing proteinase and phosphatase inhibitor cocktails, and benzonase, incubated on ice for 20 min and centrifuged for 2 min.

### Immunofluorescence staining

Cells were cultured on chamber slides (LAB-TEK) and fixed with 4% paraformaldehyde solution for 5 min and permeabilized with 0.3% Triton X-100. After washing three times with PBS and blocking with 5% BSA for 1 hr, cells were labeled with rabbit polyclonal anti-TFE3 antibody and then incubated with AlexaFluor 546 conjugated anti-rabbit antibody. Cells were covered with coverslips with Prolong Gold Antifade reagent (Invitrogen). Subcellular localization of TFE3 was examined under fluorescence microscope (Carl Zeiss).

### GST, GST-Tfe3 and GST-S6K1 protein purification

Mouse TFE3 (Tfe3) was amplified by PCR using the primers (5’-GGTTCCGCGTGGATCCATGTCTCATGCAGCCGAGCCA-3’ and 5’-GGAATTCCGGGGATCCTCAGGACTCCTCTTCAATGCTGA-3’) and cloned into BamHI site of pGEX-4T-1 expression vector (GE Healthcare) using In-Fusion HD Cloning System (Clontech). Recombinant GST and GST-Tfe3 proteins were expressed in *E.coli* [Rosetta (DE3)pLysS, Millipore] and isolated using Pierce GST Spin Purification Kit (Thermo Scientific). GST-p85-S6K1 expression vector^48^ was obtained from Addgene (Plasmid #8466) and transfected into HEK293A cells using Fugene 6 transfection reagent. Two days after transfection, 1μM mTOR inhibitor PP242 were added to culture media for 30 min prior to cell harvest. Cells (10 x 10 cm dishes) were lysed in 10 ml of CHAPS buffer (40 mM HEPES [pH 7.4], 2 mM EDTA, 10 mM pyrophosphate, 10 mM glycerophosphate, 0.3% CHAPS) supplemented with proteinase inhibitors (Roche). Cell lysates were incubated with 200μl of GST-Sepharose beads (Thermo Scientific) and washed 6 times with CHAPS buffer containing 0.5 M NaCl and 3 times with CHAPS buffer containing 5 mM MgCl_2_. GST-S6K1 proteins were eluted with storage buffer (20 mM HEPES [pH 8.0], 200 mM NaCl, and 5 mM MgCl_2_, 20% glycerol) supplemented with 10 mM glutathione.

### mTORC1 *in vitro* kinase assay

HEK293A cells were transfected with HA-Raptor and myc-mTOR expression vectors and were treated with 1 ng/ml of insulin for 30 min prior to harvest. Cells transfected with pEGFP-C2 expression vectors were used as negative control. Cells were washed once with cold PBS and lysed in CHAPS buffer containing proteinase inhibitors. Insoluble materials were removed by centrifugation for 10 min at 13,000 g. mTORC1 complex (HA-Raptor and myc-mtor) was immunoprecipitated using anti-HA antibody coupled to Protein G-magenetic beads (Dynabeads Protein G) for 2 h at 4°C. Immunoprecipitates were washed twice in low salt CHAPS buffer containing 150 mM NaCl, twice in high salt CHAPS buffer containing 400 mM NaCl and twice in HEPES/KCl buffer (25 mM HEPES pH 7.4, 20 mM KCl). mTORC1 (HA-Raptor and myc-mTOR) immunoprecipitates were incubated for 30 min at 37°C in mTORC1 assay buffer (25 mM HEPES [pH 7.4], 50 mM KCl, 10 mM MgCl_2_, 250 μM ATP and 10 μCi of ^32^P γ-ATP [3,000 Ci/mmol]) containing 60-100 ng of GST, GST-Tfe3 or GST-p85-S6K1 as substrate. Reactions were stopped by the addition of Laemmli Sample Buffer and boiling for 10 min, resolved by 4-20% or 10% SDS-PAGE. Gels were fixed with 40% methanol and 10% acetic acid for 30 min, washed three times with water, stained with colloidal Coomassie Blue (Bio-Rad), dried and exposed to X-ray film.

### AMPK *in vitro* kinase assay and TFE3 phosphorylation site analysis

AMPK (α2β1γ1, 50 ng) was incubated with GST (100 ng), GST-Tfe3 (60 ng) or SAMS peptide (1 μg) in a buffer (20 mM Tris-HCl, pH7.5, 10 mM MgCl_2_, 5 mM DTT, 240 μM ATP, 10 μCi ^32^P-γ-ATP) in the presence or absence of AMP (240 μM) for 30 min at room temperature. Reactions were terminated by adding equal volume of Laemmli Sample Buffer. After boiling for 10 min, reaction mixtures were resolved by 4-20% SDS-PAGE. Gels were fixed and stained as described above. For mass-spectrometry, GST-Tfe3 (600 ng) was phosphorylated by AMPK (500 ng) in the absence of ^32^P-γ-ATP. As a negative control, GST-Tfe3 was incubated with the same buffer but AMPK was omitted. GST-Tfe3 protein bands were excised and digested with trypsin. Peptides were analyzed by liquid chromatography-tandem mass spectrometry (MS/MS) in the Proteomics Core Facility at the University of Pennsylvania. Scaffold 3 software (Proteome Software Inc.) was used for the analysis of peptides and the phosphorylation sites.

## Results

### FLCN negatively regulates TFE3-mediated lysosomal gene expression

As reported previously^14^, a renal cancer cell line, UOK257, deficient in *FLCN* tumor suppressor gene expression, expressed two isoforms of active, hypo-phosphorylated TFE3 transcription factor (TFE3^72kDa^ and TFE3^89kDa^) (Fig 1A). Re-expression of FLCN in UOK257 cells induced phosphorylation of both TFE3 isoforms. The protein expression of TFE3 targets, such as GPNMB, ATP6V1G1 (a subunit of V-ATPase) and mucolipin-1 (a cation channel in the lysosome), were reduced upon adenovirus-mediated FLCN-expression in UOK257 cells (Fig 1B). Ectopic expression of TFE3 augmented the levels of GPNMB and ATP6V1G1 in FLCN-null UOK257 cell lysates although mucolipin-1 expression was not further increased. Similar to FLCN re-expression, *TFE3* knockdown by shRNA in FLCN-null cells decreased the expression of GPNMB (Fig 1C). Quantitative RT-PCR demonstrated that the mRNA expression of lysosomal genes, such as *acid ceramidase* (*ASAH1*), *cathepsins* (*CTSA* and *CTSB*) *lysosomal neuraminidase* (*NEU1*), *mucolipin-1* (*MCOLN1*) and mucolipin-3 (*MCOLN3*), but not mucolipin-2 (*MCOLN2*) were reduced upon TFE3 knock-down by shRNA (Fig 1D). Conversely, adenovirus-mediated overexpression of TFE3 induced lysosomal gene expression in both FLCN-null UOK257 and HEK293 cells although there appeared to be cell-type specificity (Fig 1E & 1F). Some lysosomal genes such as *ATP6V0E1* (a subunit of v-ATPase), *CTSB, CTSK* and *MCOLN3* were induced only in UOK257 but not in HEK293 cells. On the other hand, *NEU1* expression was induced only in HEK293 cells suggesting a cell-type specific role of TFE3. Importantly, overexpression of TFE3 in UOK257 cells acidified culture media as demonstrated by the pH indicator, phenol red (Fig 1G). Acidification stimulated by TFE3 was completely suppressed by the addition of a V-ATPase inhibitor, bafilomycin A suggesting involvement of V-ATPase in the acidification. Ectopic TFE3 expression not only made cells lose contacts with neighboring cells but also changed cellular morphology whereas GFP control expression did not influence UOK257 cell-cell contacts (Fig 1H). These data indicate that TFE3, which is activated by mutational inactivation of *FLCN*, induces extracellular acidification and cellular morphological changes that affect cell-cell contacts, possibly through expression of lysosomal enzymes such as cathepsins, mucolipins and V-ATPase.

**Fig 1.**
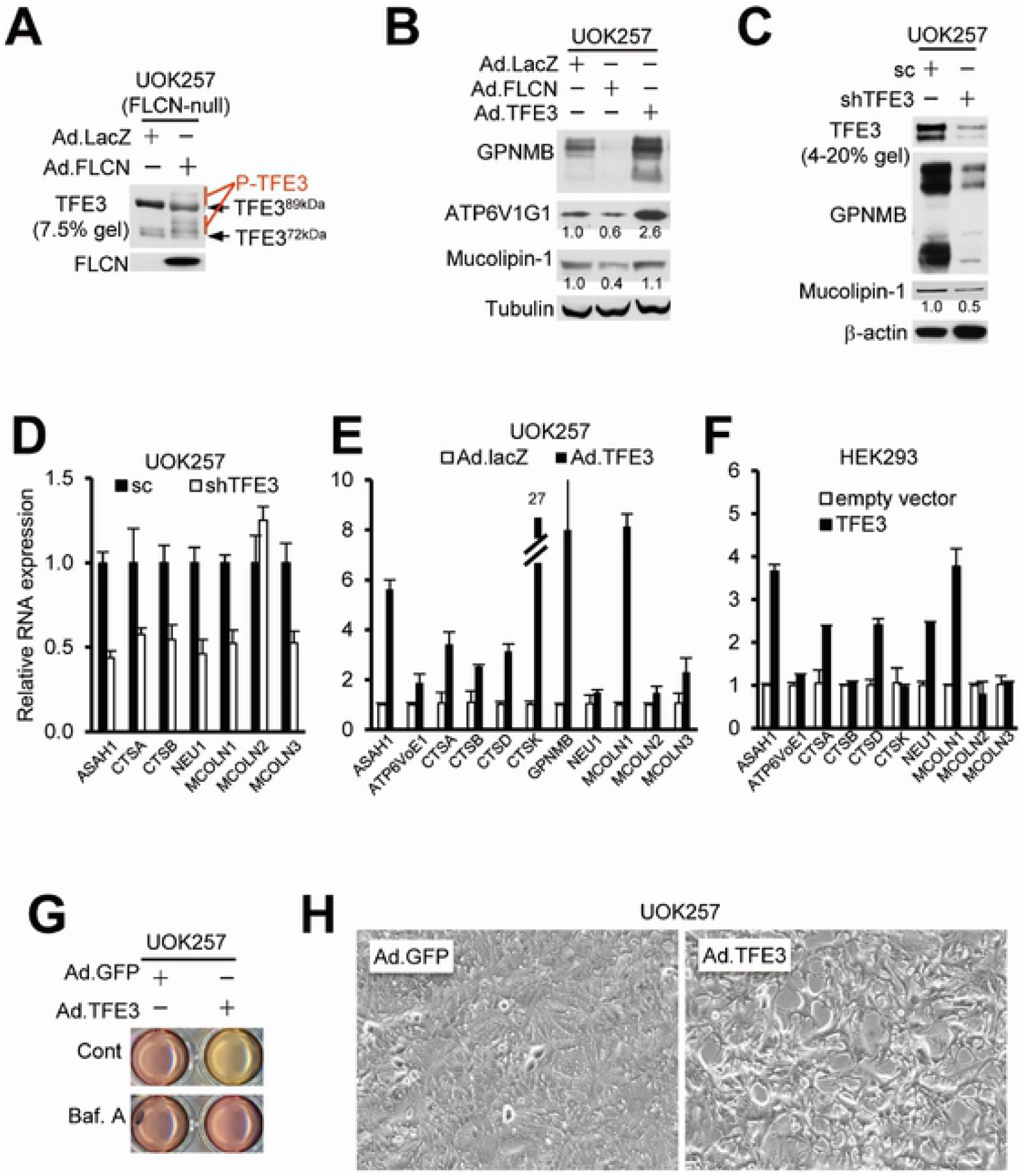
Lysosomal gene expression, acidification of media and cellular morphological changes induced by TFE3 transcription factor. (A) Increased level of TFE3 phosphorylation by the transduction of adenovirus expressing FLCN (Ad.FLCN) into FLCN-null UOK257 cells. Ad.lacZ was used as a negative control. (B) Regulation of TFE3 target expression by FLCN and TFE3. UOK257 cells were infected with Ad.lacZ, Ad.FLCN and Ad.TFE3 and were harvested at 48 hr. The protein expression of TFE3 targets in the cell lysates was measured by Western blot analysis. The expression of acid ceramidase in the culture media (media) was also measured. (C) Reduction of TFE3 and target gene (GPNMB and Mucolipin-1) expression by TFE3 shRNA in UOK257 cells. Quantitative RT-PCR of lysosomal gene expression in (D) UOK257 cells expressing either scrambled shRNA (sc) or TFE3 shRNA (shTFE3); (E) UOK257 cells expressing either Ad.lacZ or Ad.TFE3; (F) HEK293 cells transfected with either empty vector or TFE3 expression vector. Relative RNA expression and standard deviations were graphed. (G) V-ATPase-dependent acidification of culture media by TFE3. UOK257 cells were transduced with Ad.GFP or Ad.TFE3 and treated with bafilomycin A (Baf. A, 25 nM). (H) UOK257 cells were transfected with either Ad.GFP or Ad.TFE3 and their morphological changes were observed at 48 hr. All the graphs depict average values and standard deviations.

### Nutrient deprivation activates TFE3 transcription factor by inducing its nuclear localization

In order to understand the mechanism of TFE3 regulation, we investigated the physiological stimuli that affect the post-translational modification and subsequent cytoplasmic-nuclear shuttling of TFE3. Since lysosomal enzymes are involved in catabolic processes, we investigated whether metabolic stresses induce lysosomal gene expression through TFE3. TFE3 localized preferentially in the cytoplasm of mouse embryonic fibroblasts (MEF) under nutrient rich growth conditions (Fig 2A). Upon glucose deprivation, TFE3 translocated into the nucleus in about 56% of wildtype MEFs (55.8±2.1%, P<0.001). Although nuclear localization of TFE3 was also observed in AMPK-null (AMPK-KO) MEFs, it was significantly lower (~22.1±3.7%, P<0.001) than in wildtype (WT) MEFs. Fractionation experiments with HEK293 cells demonstrated that either glucose or amino acid deprivation reduced TFE3 levels in the cytoplasm by 0.5fold (P<0.05) but induced TFE3 levels in the nucleus by 30fold (P<0.01) (Fig 2B). mTORC1 activity as measured by S6K1 phosphorylation was suppressed by either glucose or amino acid deprivation (Fig 2B). On the other hand, AMPK activity was only induced by glucose deprivation. TFE3 hypo-phosphorylation in the cytoplasm and increased nuclear localization of TFE3 by glucose-deprivation were also observed in other cell lines such as NIH3T3, MDCK and HeLa cells (Fig 2C). Addition of 2-deoxyglucose (2-DG), an inhibitor of glycolysis, reduced the phosphorylation level of TFE3 and S6K1 (Fig 2D). Overall, our data demonstrated that metabolic stresses that activate AMPK and/or inactivate mTORC1 induced nuclear localization of TFE3 concomitant with its hypo-phosphorylation.

**Fig 2.**
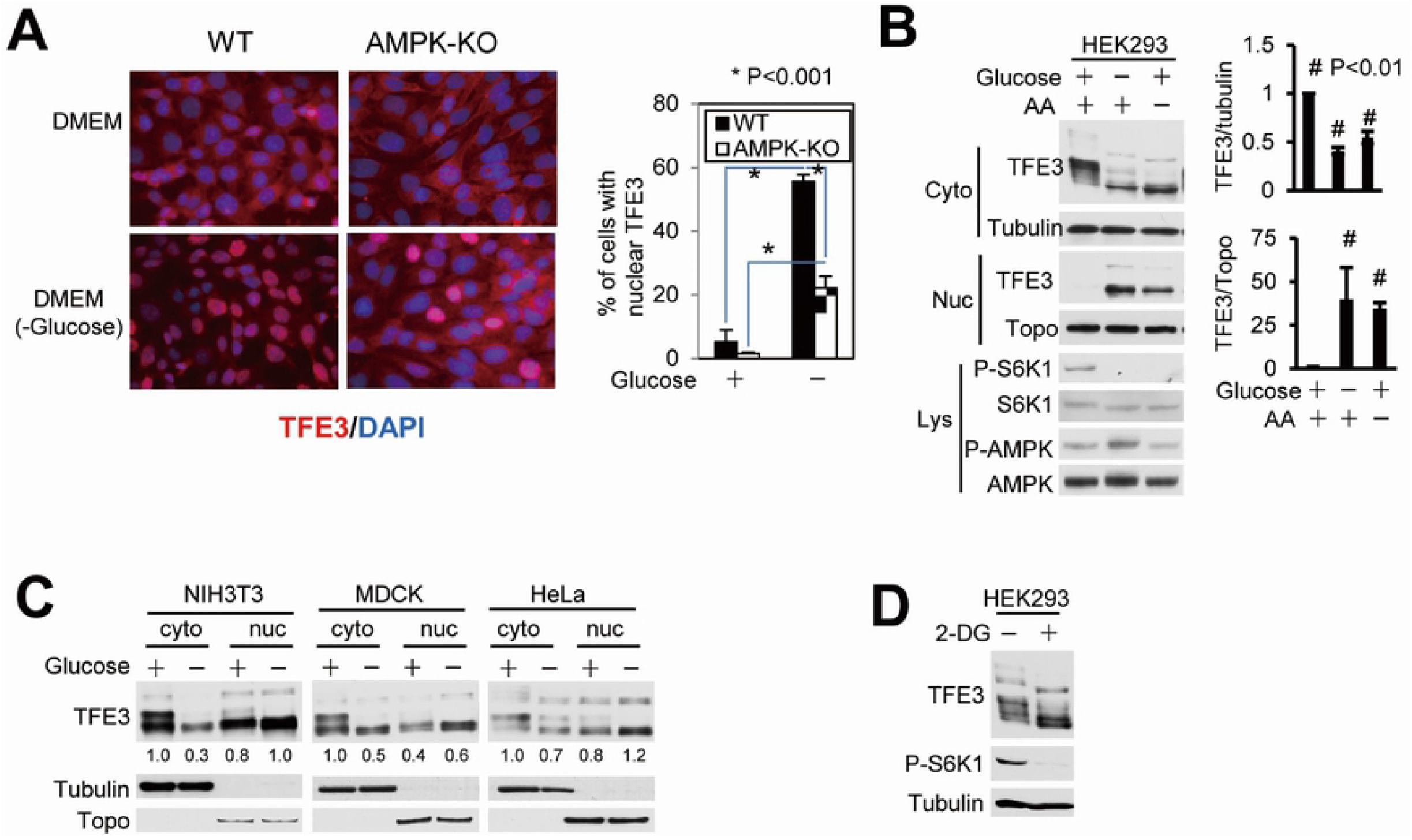
Nuclear localization and hypo-phosphorylation of TFE3 induced by glucose or amino acid deprivation. (A) Wildtype (WT) and AMPK-null (AMPK-KO) MEFs were subjected to glucose deprivation for 30 min. Subcellular localization of TFE3 was determined by immunofluorescence staining. The graph shows the percentage of cells with nuclear staining of TFE3 and the statistical analysis. N>500, *P<0.001 (student’s t-test). (B) Fractionation experiment in HEK293 cells demonstrating the reduced level of cytoplasmic (Cyto) and increased level of nuclear (Nuc) TFE3 after glucose or amino acid deprivation for 1 hr. The graphs show the level of TFE3 in the cytoplasm and nucleus normalized to tubulin and topoisomerase 1 (Topo), respectively. N=3, #P<0.05, *P<0.01 (student’s t-test). WCL, whole cell lysate. (C) Hypo-phosphorylation and nuclear localization of TFE3 in response to glucose deprivation in NIH3T3, MDCK and HeLa cells. (D) Reduced phosphorylation levels of TFE3 and S6K1 in HEK293 cells treated with a glycolytic inhibitor, 2-deoxyglucose (2-DG, 20 mM).

### AMPK activation and/or mTORC1 inhibition induced lysosomal gene expression through TFE3 transcription factor activation

We then investigated whether AMPK and mTORC1 play roles in the regulation of TFE3 phosphorylation. AICAR, a specific pharmacological activator of AMPK, induced nuclear localization of TFE3 in an AMPK-dependent manner (Fig 3A). In response to AICAR treatment, TFE3 translocated into the nucleus in AMPK-proficient wildtype MEFs but not in AMPK-null MEFs. AMPK activation by AICAR induced TFE3 hypo-phosphorylation; however, Compound C, a pharmacological inhibitor of AMPK, suppressed AICAR-induced hypo-phosphorylation of TFE3 in wildtype but not in AMPK-KO MEFs (Fig 3B). Fractionation experiments in HEK293 cells confirmed translocation of TFE3 from cytoplasm to nucleus in response to AICAR treatment (Fig 3C). In addition, mTOR inhibitors rapamycin and pp242 induced nuclear localization of TFE3. As expected, in addition to increased nuclear TFE3, TFE3-target gene expression was induced by glucose or amino acid deprivation, AMPK activator or mTOR inhibitor in the wildtype MEFs as demonstrated by elevated mRNA expression of TFE3 target genes *Gpnmb* and *Mcoln1* (Fig 3D). TFE3-target genes were also induced similarly by glucose deprivation but to a lesser degree by amino acid deprivation in AMPK-null MEFs compared with wildtype MEFs. While AICAR induced TFE3-target gene expression in an AMPK-dependent manner, mTORC1 inhibitor (pp242) induced TFE3 target gene expression even in the absence of AMPK expression. Interestingly, the levels of TFE3-target gene expression tend to be higher in wildtype MEFs compared with AMPK-null MEFs. Overall, our data demonstrated that the AMPK-mTOR signaling axis plays an important role in the regulation of TFE3 by modulating its subcellular localization (Fig 3D).

**Fig 3.**
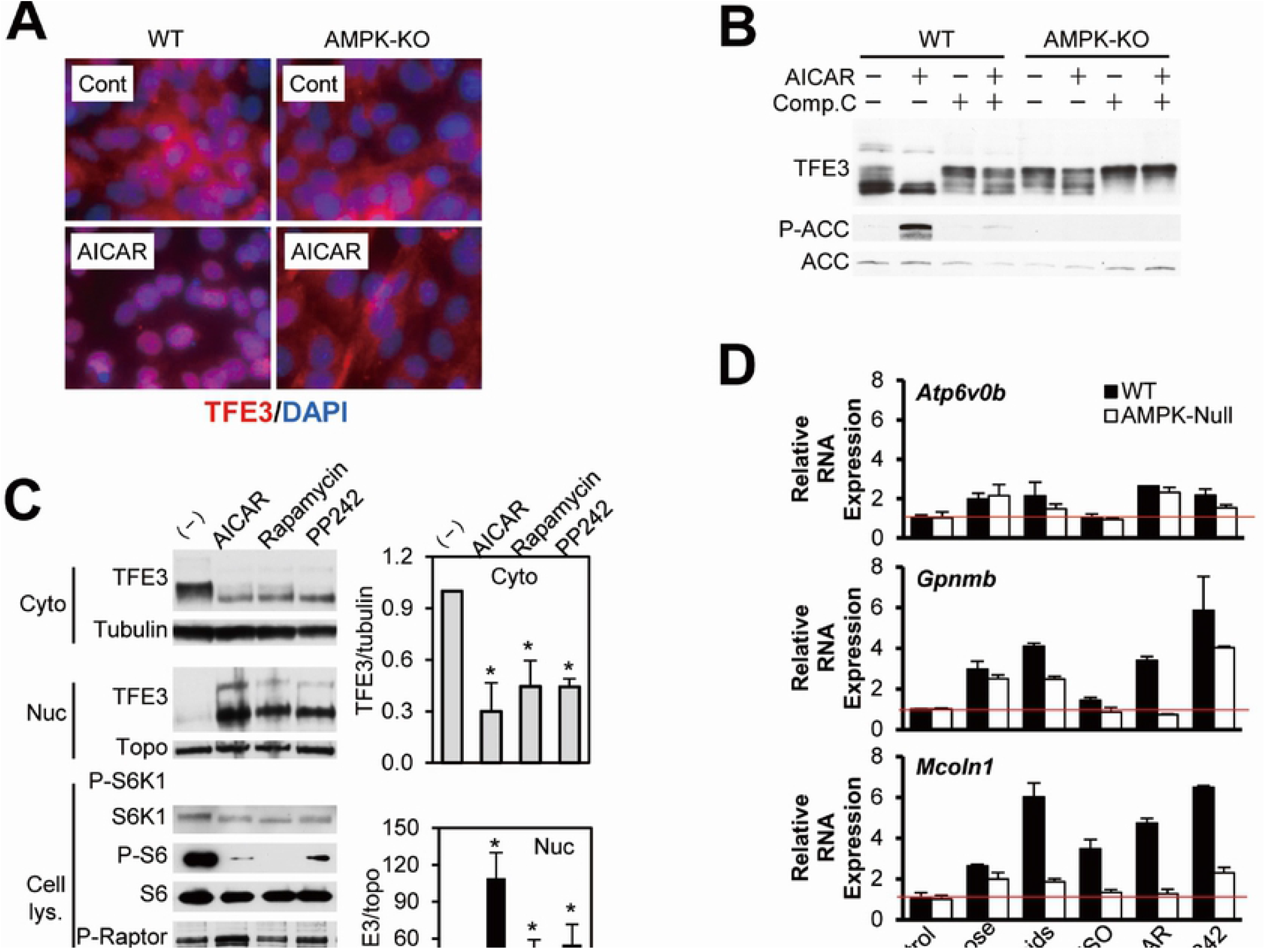
Regulation of TFE3 activity by the AMPK-mTORC1 signaling axis. (A) Representative immunofluorescence staining of TFE3 in wildtype (WT) and AMPK-null (AMPK-KO) MEFs treated with AICAR (2 mM). The graph demonstrates percentage of cells with nuclear TFE3 staining (N>300 cells each). (B) Changes in TFE3 phosphorylation in WT and AMPK-KO MEFs treated with AICAR (2 mM) and Compound C (20 μM) alone and in combination. (C) Fractionation of HEK293 cells treated with AICAR (2 mM), rapamycin (200 nM) or PP242 (200 nM) for 1 hr. The graphs show statistical analysis of the subcellular localization of TFE3. Levels of TFE3 in cytoplasm and nucleus were normalized to tubulin and topoisomerase 1, respectively. N=3, *P<0.05 (student’s t-test). (D) Quantitative RT-PCR of TFE3-target gene expression (*Gpnmb* and *Mcoln1*) in the wildtype and AMPK-KO MEFs cultured in control, glucose-free or amino acid-free media, or treated with DMSO, AICAR (2 mM) or PP242 (1 μM) for 6 hr.

### TFE3 phosphorylation by mTORC1 and AMPK

We investigated whether mTORC1 directly phosphorylates TFE3 using an *in vitro* kinase assay. Active mTORC1 was obtained from HEK293 cells transfected with HA-Raptor and myc-mTOR as described in Materials and Methods (Fig 4A). mTORC1 phosphorylated both recombinant mouse TFE3 and S6K1 fused with GST but did not phosphorylate GST alone (Fig 4A). Phosphorylation of TFE3 and S6K1 by mTORC1 was suppressed by a specific inhibitor of mTOR, PP242, confirming the specificity of TFE3 phosphorylation by mTORC1 (Fig 4A).

**Fig 4.**
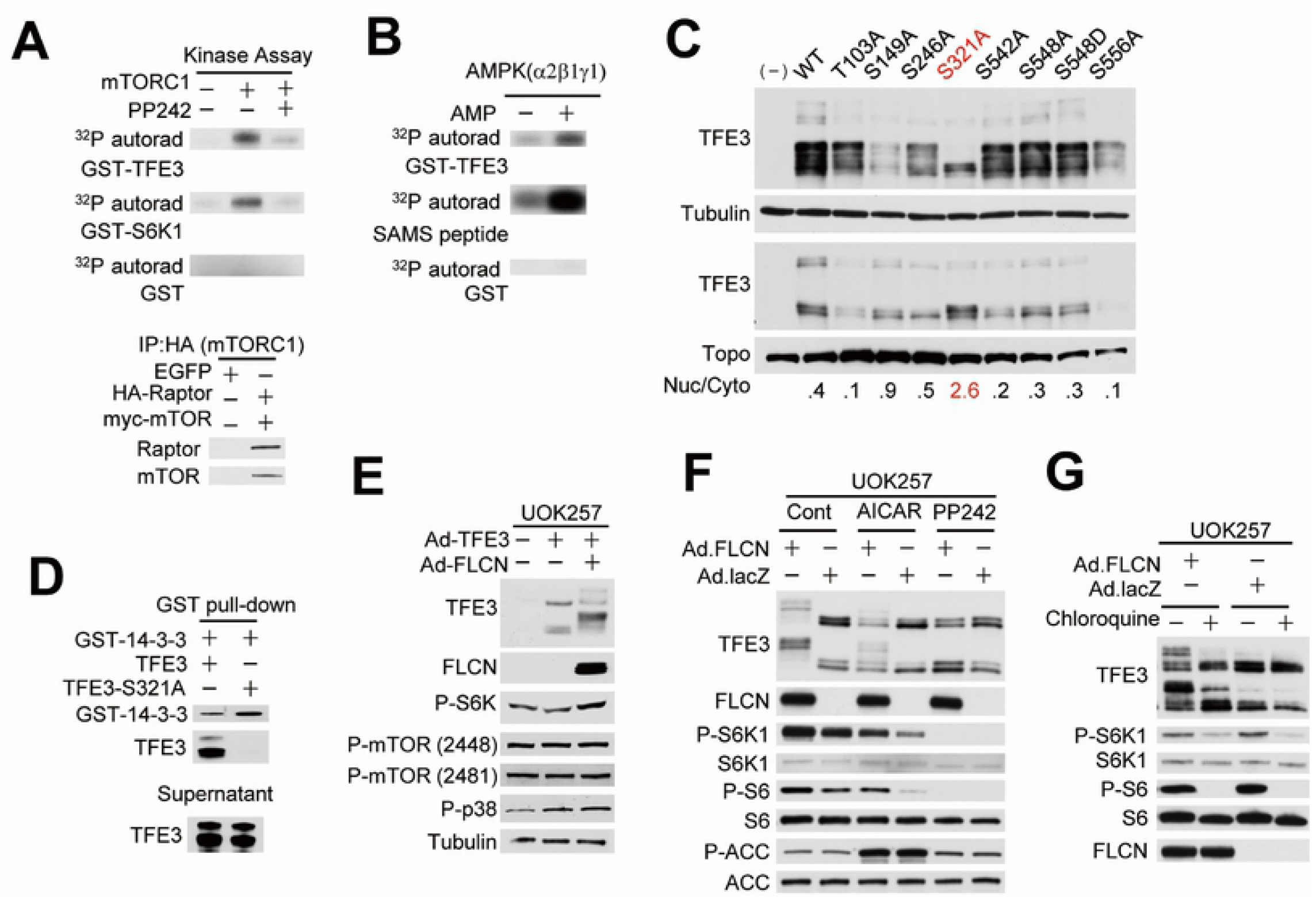
TFE3 phosphorylation by mTORC1 and AMPK, and requirement of FLCN expression. (A) TFE3 phosphorylation by mTORC1 *in vitro*. ^32^P-phosphorylation of GST-Tfe3 and GST-S6K1 but not GST by the immunoprecipitated mTORC1 kinase complex (HA-Raptor/myc-mTOR). PP242, 200 nM. IP, immunoprecipitates. (B) TFE3 phosphorylation by AMPK *in vitro*. GST-TFE3, SAMS peptide and GST were incubated with recombinant AMPK (α2β1γ1) in the presence or absence of AMP (400 μM). (C) Fractionation of HEK293 cells transfected with wildtype or mutant TFE3 expression vectors. The ratios of nuclear to cytoplasmic TFE3 (Nuc/Cyto) are indicated. (D) GST pull-down experiment demonstrating the interaction between wildtype TFE3 (but not TFE3-S321A mutant) and 14-3-3 protein. (E) FLCN expression was required for TFE3 phosphorylation in cells under nutrient-rich conditions. Both mTORC1 activity and FLCN expression were required for TFE3 phosphorylation. UOK257 cells expressing either FLCN or lacZ were treated with (F) DMSO (Cont), AICAR (2 mM), PP242 (1 μM) and (G) chloroquine (100 μM). (H) Schematic diagram demonstrating a proposed mechanism for TFE3 regulation by the FLCN-AMPK-mTORC1 signaling network.

We also investigated whether TFE3 is phosphorylated by AMPK directly. Interestingly, recombinant AMPK (α2β1γ1), in the presence of AMP, phosphorylated recombinant GST-Tfe3 protein and SAMS peptide, a well-known substrate of AMPK, but not GST protein itself (Fig 4B). We then determined the AMPK-mediated TFE3 phosphorylation sites by mass spectrometry as described in the Materials and Methods. Mass spectrometry covered only 159 out of 537 amino acids (30%) and identified 10 phosphorylated Serine or Threonine residues in Tfe3 (STable 1). A peptide containing a phosphorylated serine residue at codon 510 (S510) in mouse Tfe3 isoform 4 (corresponding to S548 in human TFE3) was found most often (13/26 peptides, 50%).

In order to determine the critical residues that are important for the subcellular localization of TFE3, we predicted mTORC1-dependent phosphorylation sites using the position-specific scoring matrix for the consensus mTOR phosphorylation motif^49^. We selected 6 putative mTORC1-dependent phosphorylation sites (T103, S149, S246, S321, S542, and S556) and an AMPK-dependent phosphorylation site (S548) for analysis. We generated phosphorylation defective mutants for the selected residues by mutating serine or threonine to alanine. A phospho-mimic mutant was also generated for S548 by mutating serine to aspartic acid. The mammalian expression vectors for wildtype and mutant TFE3 were transfected to HEK293 cells and their phosphorylation status and subcellular localization were analyzed. Among the mutants, only the TFE3-S321A mutant was hypo-phosphorylated in the cytoplasm and showed a 6-fold higher nuclear to cytoplasmic TFE3 ratio compared with wildtype TFE3 (Fig 4C). It is noteworthy that S321 is in proximity to a putative nuclear localization signal (NLS) within the basic-helix-loop-helix-leucine-zipper (bHLH-LZ) domain and is conserved in the MiTF/TFE family as well as in TFE3 orthologues (S1 Fig A-C). The mutations in S548 (both S548A and S548D) did not alter nuclear to cytoplasmic TFE3 ratio. Since 14-3-3 protein has been shown to be involved in the cytoplasmic localization of MiTF/TFE transcription factors^16,27,29^, we examined whether S321 in TFE3 is important for the interaction between TFE3 and 14-3-3 protein by using a GST pull-down assay. Wildtype TFE3 (hyper-phosphorylated) interacted but the TFE3-S321A mutant (hypo-phosphorylated) did not interact with 14-3-3 protein (Fig 4D). Taken together these data indicate that phosphorylation of TFE3 at S321 by mTORC1 induces its cytoplasmic localization through association with 14-3-3 proteins.

### TFE3 phosphorylation requires both mTORC1 activity and FLCN expression

TFE3 phosphorylation was dependent on FLCN expression under nutrient and growth factor rich conditions, in which mTORC1 activity, as measured by readouts P-S6K1 (T389), P-mTOR (S2448) and P-mTOR (S2481), was similarly high in UOK257 cells (FLCN-null) expressing lacZ or re-expressing FLCN (Fig 4E). With FLCN re-expression, the level of TFE3 phosphorylation was dependent on mTORC1 activity (Fig 4F). Partial or complete suppression of mTORC1 activity by AMPK activator AICAR or mTOR inhibitor PP242 reduced TFE3 phosphorylation accordingly. In addition, chloroquine, a lysosomal inhibitor that suppresses mTORC1 activity, reduced the level of TFE3 phosphorylation in FLCN-proficient cells (Fig 4G). On the other hand, TFE3 proteins were maintained in their hypo-phosphorylated forms irrespective of mTORC1 activity in FLCN-null cells (Fig 4E, 4F and 4G). Taken together, these results indicate that FLCN expression is required for mTORC1-mediated phosphorylation of TFE3.

## Discussion

Our study demonstrates that the oncogenic transcription factor TFE3 is activated by metabolic stresses through AMPK-mTOR signaling pathway. Metabolic stresses such as glucose or amino acid deprivation or the glycolysis inhibitor, 2-DG, activated TFE3 through post-translational modifications and subsequent nuclear localization. Activation of AMPK and/or inhibition of mTORC1 activity by metabolic stresses induced expression of TFE3 targets including a number of lysosomal genes. *In vitro* kinase assays demonstrated that AMPK and mTORC1 directly phosphorylate TFE3. It is a novel finding that TFE3 can be phosphorylated by AMPK although the functional significance has not been demonstrated in this study. However, TFE3 phosphorylation by mTORC1 and subsequent interaction with 14-3-3 protein is a likely mechanism for the cytoplasmic retention of TFE3. FLCN expression was essentially required for phosphorylation by mTORC1 and cytoplasmic retention of TFE3.

Here we demonstrated the importance of S321 residue in TFE3, a putative mTORC1-mediated phosphorylation site, in the interaction with 14-3-3 protein and cytoplasmic localization. Similarly, to our results, Martina et al. demonstrated mTORC1-mediated phosphorylation of S321 in TFE3 and its role in cytoplasmic localization^16^. It was also reported that TFEB, a closely related transcription factor, is negatively regulated by mTORC1^16,21,30^. S211 in TFEB, which is equivalent to S321 in TFE3, is an mTORC1-dependent phosphorylation site that is important for the interaction with 14-3-3 protein and cytoplasmic localization of TFEB. The subcellular localization of MiTF is also regulated by a similar mechanism. Phosphorylation of MiTF on S176, which is equivalent to S321 in TFE3, is required for interaction with 14-3-3 protein and the cytoplasmic localization of MiTF^28,33^. Accordingly, it is likely that the subcellular localization of the members of the MiTF/TFE transcription factor family is regulated by phosphorylation and interaction with 14-3-3 protein.

The lysosome is an important subcellular compartment for catabolic processes but also for metabolic signaling. mTORC1 is translocated to the lysosomal membrane and activated in the presence of amino acids^3^. V-ATPase activity, which acidifies the lysosomal lumen, plays an important role in promoting mTORC1 translocation to the lysosome and its activation^4^. Inhibitors of V-ATPase and lysosomal function, such as salinomycin A and chloroquine, suppress mTORC1 activity. Previous reports and our current study demonstrate the concomitant reduction of TFEB and TFE3 phosphorylation by mTORC1 inhibition^4,20,30^. The resulting activation of TFE3 and TFEB induces lysosomal function through the expression of lysosomal genes.

The functional roles of lysosomal enzymes in the pathogenesis of human disease including cancer have been elucidated to a great extent^11^. Cathepsins are lysosomal proteinases that are highly up-regulated in a wide variety of cancers^50–52^. Although cathepsins are localized within intracellular lysosomes, their localization is often altered during neoplastic progression resulting in their secretion. Upregulation and altered localization of cathepsins induce tumor cell growth, migration, invasion, angiogenesis and metastasis. Recently it was demonstrated that a lysosomal cation channel, mucolipin 1, a transcriptional target of TFEB and TFE3, induces exocytosis of lysosomal hydrolases by releasing calcium from the lysosomal lumen into the cytosol^17^. Lysosomal exocytosis also alters the subcellular localization of the lysosomal proton pump V-ATPase to the plasma membrane^53^. V-ATPase, localized in the plasma membrane, pumps out protons into the extracellular space. The resulting acidic micro-environment will activate secreted lysosomal hydrolases and thus promote degradation of extracellular matrices, a process that is necessary for tumor progression^54^. Overexpression of TFE3 induces the acidification of culture media in a V-ATPase-dependent manner and disrupts cell-cell contacts as shown in Fig 1G&H. Our data suggest that TFE3 and its family of transcription factors play important roles in the pathogenesis of human disease through enhancing the lysosomal enzyme activities in the lysosome, plasma membrane and peri-cellular spaces.

In this study, we showed that AMPK phosphorylates TFE3 *in vitro*. We examined whether the mutations in S548, the most prominently AMPK-dependent phosphorylation site, would change the profile of TFE3 subcellular localization. Contrary to our expectation, the subcellular localization of TFE3 was not affected by the mutations in S548. We cannot rule out the possibility that other untested AMPK-dependent phosphorylation sites might play roles in TFE3 subcellular localization. On the contrary, AMPK might play a role in fine tuning TFE3 activity through augmentation of its transcriptional activity, retention in nucleus or modulating target selection. In support of this idea, the gene expression data in Fig 3D show that mucolipin-1 expression was induced by metabolic stresses to a much higher level in wildtype (AMPK-proficient) cells compared with AMPK-KO cells. The functional roles of AMPK in the regulation of transcription have been demonstrated. AMPK directly phosphorylates several transcription factors and cofactors including SREBP1, ChREBP and CRTC2, all of which are involved in the control of metabolic gene expression. It is of interest that AMPK inhibits SREBP1, which competes with TFE3 and FOXO1 complex for binding to IRS2 promoter^55^. Further analysis analyzing the physiological significance of TFE3 phosphorylation by AMPK would be necessary.

Recently, a pro-tumorigenic role of AMPK-TFE3 axis was demonstrated in Kras-dependent non-small-cell cancer model^44^. AMPK was required for growth of lung tumors with oncogenic Kras mutation (Kras^G12D^). In addition, AMPK played a critical role in regulating lysosomal gene expression through Tfe3, which was required for tumor growth. In addition, AMPK-dependent regulation of TFEB has also been demonstrated. AMPK played critical roles in lineage specification through TFEB-dependent regulation of lysosome^56^. Whole-genome transcriptome profiling in wild-type and AMPK-KO cells revealed that not only lysosomal genes but also the FLCN gene was regulated by AMPK through TFEB^57^. Contrary to the current study, AMPK promoted dephosphorylation and nuclear localization of TFEB independent of mTORC1 activity. This suggests that AMPK plays multiple roles in the regulation of TFEB dephosphorylation other than inhibition of mTORC1.

How does loss of FLCN activate TFE3? It has been demonstrated that not only TFEB but also TFE3 is recruited to lysosome through interaction with active Rag GTPases and is thereby phosphorylated by mTORC1 in a nutrient replete condition^58–59^. FLCN is also recruited to lysosome by binding to Rag GTPases and acts as a guanine nucleotide activating protein (GAP) for the RagC/D GTPase, which facilitates Rag GTPases to associate with mTORC1 and thus leads to mTORC1 activation^41,60^. Thus loss of FLCN is likely to maintain Rag GTPases in an inactive state, which would block recruitment of mTORC1 and TFE3 on the lysosome and thus inhibit phosphorylation of TFE3 by mTORC1. Although loss of FLCN inhibited mTORC1 activity toward TFE3 phosphorylation, mTORC1 activity toward S6K1 remained intact in nutrient- and growth factor-enriched condition. Unbalanced regulation of mTORC1 activity, which on the one hand induces catabolic processes through activation of TFE3, while supporting anabolic processes through activation of S6K1, is likely to contribute to tumorigenesis.

Another possibility is that elevated expression of mucolipin-1, a lysosomal calcium channel contributes to the induction of TFE3 dephosphorylation by calcineurin and subsequent nuclear localization. Dephosphorylation of TFEB is mediated by the phosphatase calcineurin (CaN), which is activated following mucolipin-1-mediated lysosomal calcium release^61^ and enables TFEB to translocate into the nucleus^61^. A similar mechanism may be regulating the subcellular localization of TFE3 following AMPK activation and/or loss of FLCN.

TFE3 immunoreactivity has been demonstrated in 9% of all renal carcinomas and is associated with high nuclear grade, greater tumor extent of spread, metastasis and poor patient outcome^27^. However, Xp11.2 translocations involving TFE3 have been identified in less than 1% of all renal carcinomas. Accordingly, other genetic changes including loss of *FLCN* or *TFE3* gene amplification, might account for the remainder of renal cancers. Elevated lysosomal gene expression during the progression of various cancers may be in part due to the activation of TFE3 or its related transcription factors. Activation of TFE3 has been shown in a subset of renal cell carcinoma and alveolar soft part sarcoma. However, we cannot exclude the possibility that TFE3 may play a role in more diverse types of cancers. Further investigations will reveal the significance of TFE3 more globally in the development and progression of cancer.

Here we have investigated the post-translational regulation of TFE3 transcription factor in the context of energy and metabolic signaling. Upon nutrient deprivation, TFE3 is activated and induces lysosomal gene expression to facilitate catabolic processes that will generate nutrients to accommodate cellular energy needs. Activation of AMPK and/or inhibition of mTORC1 activity by metabolic stresses induced nuclear localization of TFE3 and its activation. Suppression of mTORC1-mediated phosphorylation of TFE3 and maintenance of its unphosphorylated status by the inactivation of *FLCN* are likely mechanisms of TFE3 activation. Uncontrolled activation of TFE3 is likely to play an important role in the development of human disease including cancer. This study sheds light on our understanding of the mechanisms of TFE3 regulation. Further investigation will be necessary to delineate the precise mechanism by which TFE3 is being regulated by mTORC1 and AMPK.

## Acknowledgments

We thank Dr. Benoit Viollet at INSERM for providing wildtype and AMPK-null MEFs.

## Supporting Information

**S1 Fig 1. The structure of TFE3 and conserved phosphorylation sites.**

(A) The mutated residues on the schematic diagram of TFE3. (B) S321 in TFE3 (red asterisk), a 14-3-3 binding motif and the putative nuclear localization signal (NLS) are conserved among the MiTF/TFE family. (C) *, conserved phosphorylation site.

**S2 Fig. Mouse TFE3 sequences identified by Mass-Spectrometry.**

(A) Un-treated Tfe3; (B) phosphorylated Tfe3 by AMPKs. The recovered regions are highlighted by yellow; phosphorylated residues are highlighted by pink.

**S1 Table 1. The AMPK phosphorylation sites on Tfe3 identified by Mass-Spectrometry.**

The phosphorylated residues are colored in red.

## Author Contributions

Conceived and designed the experiments: SH, HL, VK, LS and WL. Performed the experiments: SH, HL and HO. Wrote and edited the paper: SH, HL, VK, BV and LS.

